# Linguistically deprived children: meta-analysis of published research underlines the importance of early syntactic language use for normal brain development

**DOI:** 10.1101/166538

**Authors:** Andrey Vyshedskiy, Mahapatra Shreyas, Rita Dunn

**Author notes:** **Abbreviations:** NOB score = number of objects score PCT score = posterior cortex territory score.

## Abstract

We analyzed all published reports of individuals not exposed to syntactic language until puberty: two feral children, who grew up without hearing any language, and eight deaf linguistic isolates, who grew up communicating to their families using homesign or kitchensign, a system of gestures which allows them to communicate simple commands but lacks much in the way of syntax. A common observation in these individuals is the lifelong difficulty understanding syntax and spatial prepositions, even after many years of rehabilitation. This debilitating condition stands in stark contrast to linguistic isolates’ performance on memory as well as semantic tests: they could easily remember hundreds of newly learned words and identify previously seen objects by name. The lack of syntactic language comprehension in linguistic isolates may stem from inability to understand words and/or grammar or inability to mentally synthesize known objects into novel configurations. We have previously shown that purposeful construction of novel mental images is the function of the lateral prefrontal cortex (LPFC) ability to dynamically control posterior cortex neurons ^1^. Here we have ranked all tests performed on linguistic isolates by their reliance on the LPFC control of the posterior cortex: a) the amount of posterior cortex territory that needs to be recruited by the LPFC and b) the number of disparate objects that have to be combined together by the LPFC in order to answer the test question. According to our analysis, linguistic isolates performed well in all tests that did not involve the LPFC control of the posterior cortex, showed decreasing scores in tests that involved greater recruitment of the posterior cortex by the LPFC, and failed in tests that involved greatest recruitment of posterior cortex necessary for mental synthesis of multiple objects. This pattern is consistent with inadequate frontoposterior connections in linguistic isolates. We discuss implications of these findings for the importance of early syntactic language exposure in formation of frontoposterior connections.

## Introduction

There is general consensus that early time-sensitive exposure to syntactic language is important for acquisition of the full extent of a complex syntactic language ^2–5^. The mechanism of this process, however, remains unclear. If the language provides children with the necessary tools to train their cognition, then the lack of language use during childhood must be associated with cognitive deficit. Studying cognitive processes in adults and adolescents who have been linguistically isolated during their childhood could therefore contribute to this discussion. These linguistically isolated individuals can be categorized into two groups by the nature of their isolation. Completely linguistically isolated children grow up with no verbal or sign communication to other humans. Syntactically isolated children, on the other hand, grow up communicating to other humans using simple signs void of syntax, grammar and spatial prepositions.

The best studied case of a completely linguistically isolated child is that of Genie (real name: Susan M. Wiley) who was isolated starting at 20 months of age until she was rescued at the age of 13 years 7 months. She was locked inside a bedroom in Los Angeles, strapped to a child's toilet during the day and bound inside a crib with her arms and legs immobilized on most nights ^6,7^. She was not allowed to vocalize, was not spoken to, could not hear family conversation, or any other language occurring in her home other than swearing (there was no TV or radio in the home). Genie emerged from isolation with no development of spoken language and no comprehension of any language with exception of a few isolated words. After Genie was rescued, she was the focus of intensive language instruction.

Genie's case has been extensively compared to that of Victor of Aveyron, an eighteenth-century French child whose life similarly became a case study for research in delayed first-language acquisition. Victor of Aveyron was a feral child who apparently lived much of his childhood alone in the woods before being found wandering the woods near Saint-Sernin-sur-Rance in France. His case was taken up by a young physician, Jean Marc Gaspard Itard, who worked with the boy for five years. Itard named the wild child Victor, and estimated that the boy was 13 years old when he initiated the study. Itard, who was interested in determining what Victor could learn, devised procedures to teach the boy words and recorded his progress ^8^.

Syntactic isolation of children stems from various causes and may occur even in a loving and cohesive family. About 90% of all congenitally deaf children are born to hearing parents ^9^. In the US, these children typically receive special services. In Latin American countries, however, deaf children may never be exposed to a formal syntactic sign language. To communicate, families usually spontaneously invent homesign (a.k.a. kitchensign), a system of iconic gestures that consists of simple signs but lacks much in the way of syntax. Note that unlike Genie and Victor of Aveyron who grew up with no language and no communication, deaf linguistic isolates are using signs to communicate to their family from the very early age. The homesign system though is lacking syntax, spatial prepositions, and other recursive elements of a formal sign language. In other words, deaf linguistic isolates grow up exposed to language, but not to syntactic language.

Deaf children who grow up using homesign for communication must be distinguished from deaf children developing in a community of other deaf children, as they are known to be able to independently invent a syntactic sign language of their own. In 1980, following the Sandinista revolution, the Nicaraguan government opened several vocational schools for deaf children. By 1983 there were over 400 students in the two schools. The school program emphasized spoken Spanish and lip reading, and discouraged the use of signs by teachers. The program failed and students were unable to learn the Spanish language in such a manner. However, the school provided fertile ground for deaf students to communicate with each other. In this process, children gradually spontaneously generated a new sign language, complete with *syntax*, verb agreement and other conventions of grammar ^10–13^. Studying generational differences between Nicaraguan children who grew up when the sign language was in its initial stage of development and those who grew up a decade later exposed to a richer vocabulary and more complex recursive elements demonstrated clear cognitive differences between the different cohorts of children ^14–16^.

Syntactically isolated children may also include those who are exposed to syntactic language but do not use it either for external or for internal communications. For the most part this group is formed by children with ASD. The primary problem in these children may be neurological, leading to a secondary problem of significant language delay. In turn, lack of syntactic language use during the critical period may result in the tertiary cognitive problems. About two thirds of children with ASD eventually demonstrate significant cognitive impairment ^17^. While the origin of cognitive impairment could stem from the primary neurological problems, it can be exacerbated by the lack of syntactic language use in childhood.

Additional evidence for the importance of early language exposure for normal cognitive development comes from the controlled randomized study of the orphaned Romanian children ^18^. When Nicolae Ceausescu banned birth control and abortion in 1966 to increase Romania's population, many overwhelmed parents left children in state institutions. By 1989 this social experiment led to more than 170,000 children living in these facilities^18^. The institutionalized toddlers did not receive much personal attention and were discouraged from vocalizing. In many instances, these toddlers grew up in environments void of any rich linguistic stimuli. In the 1990s a group of 136 toddlers free of neurological, genetic and other birth defects were randomized and half of them were assigned to a foster care, while the other half remained in such an institution. Because the toddlers were randomly assigned to foster care or to remain in an institution, unlike previous studies, it was possible to show that any differences in development or behavior between the two groups could be attributed to where they were reared^18^.

In this manuscript we initially focus on the most definitive cases of language isolation: children who have been linguistically isolated until puberty and then subjected to rehabilitation attempts for several years in their adolescence or adulthood. These cases of feral children and deaf linguistic isolates present the best opportunity to delineate the effect of syntactic language deprivation on the developing brain. In the discussion, we compare our findings to other children with relatively shorter first syntactic language deprivation periods.

## Methods

We performed PubMed and GoogleScholar search in May of 2017 for all reports of linguistic isolates. The search yielded reports of ten individuals not exposed to a syntactic language until puberty, which were subsequently tested as adults following several years of focused rehabilitation: two reports of feral children and eight reports of deaf linguistic isolates. Performance of these individuals in all kinds of linguistic and nonverbal tests is reviewed in this manuscript in terms of their PFC ability to manipulate and control their imaginary visual experience.

### Assessment of linguistic test questions

A panel of three neuroscientists classified all language comprehension and nonverbal tasks administered to linguistic isolates according to the minimal neurobiological requirements necessary to obtain the correct answer ^1^. Specifically, panel members analyzed (1) The minimum number of disparate objects that needed to be purposefully imagined together in order to answer correctly, called the number of objects (NOB) score, and (2) The type of object modification required by the question (see details below). Members of the panel analyzed each question independently of each other and then discussed questions one by one to reach a unanimous opinion.

(2.1) Establishment of the type of object modification:

For any object in the mind's eye, we can voluntarily change its color, size, position in space or rotation. The mechanisms of these processes involve PFC-controlled modification of the object's representation in the posterior cortex. As related to the posterior cortex neuronal territory, such mechanisms can be classified into three classes: 1) those that do not involve any object modification; 2) those that involve modification of activity in only the ventral visual cortex, Fig. 1; and 3) those that involve coordination of activity in both the dorsal and ventral visual cortices. For example, object comparisons does not involve modification of any object; color or size modification of an object recalled in memory is limited to the ventral visual cortex ^19^; finally, location in space modification or rotation involve coordination of activity in both the ventral and dorsal visual cortices ^20–24^. Accordingly, the three types of questions were assigned the posterior cortex territory score (PCT score) ranging from zero to two, with zero corresponding to minimal posterior cortex territory and two corresponding to greatest amount of posterior cortex territory.

**Figure 1.**
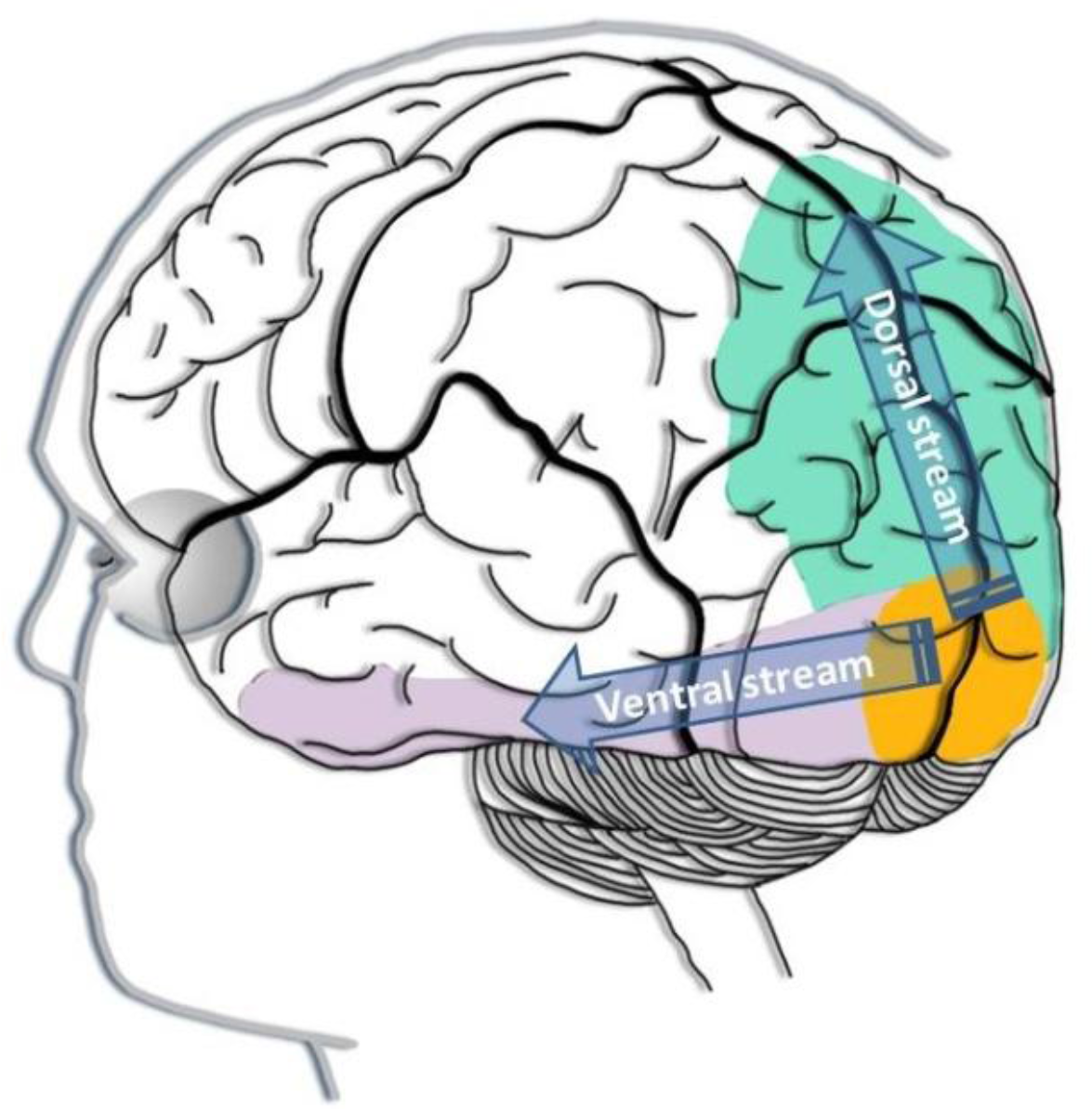
Visual information processing in the cortex. From the primary visual cortex (V1, shown in yellow), the visual information is passed in two streams. The neurons along the ventral stream also known as the ventral visual cortex (shown in purple) are primarily concerned with *what* the object is. The ventral visual stream runs into the inferior temporal lobe. The neurons along the dorsal stream also known the dorsal visual cortex (shown in green) are primarily concerned with *where* the object is. The dorsal visual stream runs into the parietal lobe.

### Language comprehension tests

#### 1. Find the named object

A common task requires the subject to find a named object from a variety of physical objects (or pictures) to choose from. These tasks are based of subject's ability to compare the internal representation of the target object with all the possible solution objects, primarily relying on matching from memory. Since this type of question does not require any mental combination of disparate objects, it is assigned a NOB score of one. Furthermore, since this type of question does not require any modification of the object, it corresponds to the PCT score of zero (i.e. requiring the least amount of the posterior cortex territory).

#### 2. Integration of modifiers in a single object

Another common comprehension test requires the subject to integrate a noun and an adjective. A subject may be asked to point out the picture with a {yellow/red/green} + circle placed among several decoy images with {yellow/red/green} + {triangle/square} thus forcing the integration of color and noun. Similarly, to integrate size and noun one may be asked to point to a {big/small} + {circle/triangle/square}. Neurologically, the integration of modifiers involves the modification of neurons encoding a *single* object and therefore has a NOB score of one. As color and/or size were modified, these tasks were given a PCT score of one. (modification is limited to the ventral visual cortex, Ref. ^19^).

##### 2.1. Comparative

In a comparative test the subject may be shown two circles of different sizes and asked to point out the circle that is {bigger/smaller}. In terms of visual representation in the posterior cortex, the circles can be considered one at a time, making the comparative task a special case of integration of the size modifier task. Therefore, minimally, the comparative task involves the modification of neurons encoding a *single* object (the circle) and has a NOB score of one. In terms of the PCT, these questions were assigned a score of one (modification is limited to the category of size represented in the ventral visual cortex, Ref. ^19^).

##### 2.2 Superlative

In a superlative test, the subjects may be shown several circles of different sizes not aligned by size and asked to point out the biggest/smallest circle. Similar to comparative task, superlative task is a special case of integration of the size modifier, it involves the modification of neurons encoding a *single* object and therefore has a NOB score of one. In terms of the PCT, these questions were assigned a score of one (modification is limited to the ventral visual cortex, Ref. ^19^).

#### 3. Integration of number modifier

To integrate the number and noun the subject may be asked to point out a card with {two/three/four} circles among distractor cards with {two/three/four} + {squares/triangles}. Similar to comparative task, the integration of number modifier minimally involves the modification of neurons encoding the number: independently of whether the number is two or a million, only a *single* object (the circle) is visualized and therefore this test has a NOB score of one. Reports indicate that numerical information is represented by regions of the posterior parietal lobes ^25,26^. Thus, these tasks are given a PCT score of two.

##### 3.1 Singular vs. plural

Singular vs. plural tasks involve showing several pictures and asking the subject to point out the correct picture: ‘Show me the flower’ versus ‘Show me the flowers. Neurologically, in terms of visual representation of object in the posterior cortex, these tasks are the special case of the integration of number modifier; they involve the modification of neurons encoding a *single* object (NOB score of one) and have a PCT score of two.

##### 3.2 Negative vs. affirmative statements

To test understanding of simple negation the subject is shown a pair of pictures identical except for the presence or absence of some element. For example, the subject may be asked to indicate a picture in which the rabbit has a carrot vs. the one in which the rabbit does *not* have a carrot. Neurologically, in terms of visual representation of object in the posterior cortex, these tasks involve the modification of neurons encoding a *single* object (e.g., a carrot) and therefore have a NOB score of one. These questions are a special case of integration of number modifier, with the number set to zero. Accordingly, they were assigned a PCT score of two.

#### 4. Mental rotation and modification of a single object's location in space

A number of verbal and nonverbal tasks given to linguistic isolates require mental rotation or other modification of an object's location in space. An example of a nonverbal mental rotation task is presented in Figure 2. Mental rotation and modification of a single object's location in space involve the PFC-coordinated activity in *both* the ventral and the dorsal visual cortices ^23,24,27^, and are therefore assigned the PCT score of two. However, these tasks are still limited to a single object and, consequently, assigned a NOB score of one.

**Figure 2.**
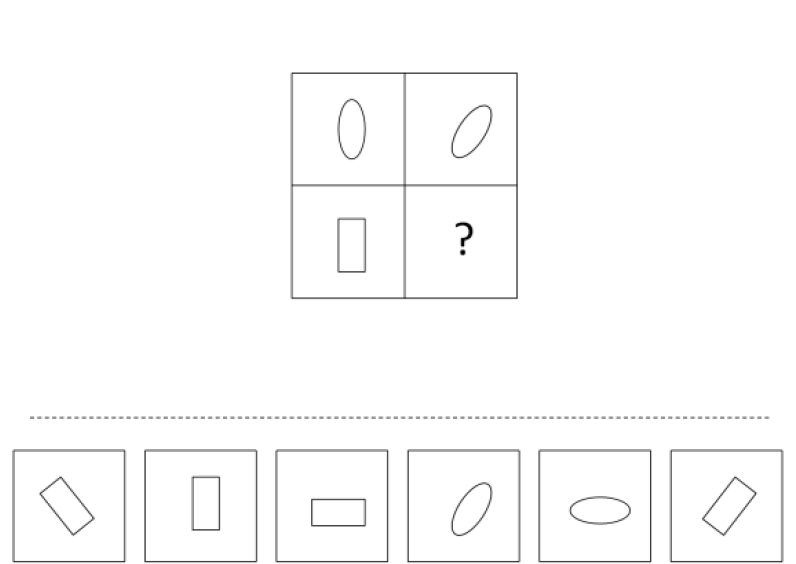
A typical question testing subject's ability to mentally rotate an object is shown here as a 2×2 matrix with six answer choices displayed below the problem. The top row of the matrix indicates the rule: “the object in the right column is the result of 45**°** clockwise rotation.” Applying this rule to the bottom row, we arrive at the correct answer depicted on the right.

##### 4.1 Copying stick and block structures

Copying stick and block structures test is a nonverbal test that relies on mental rotation of sticks or block pieces in order to align them with the instructions. Since mental manipulations involve one piece at a time, these tasks are limited to a single object and, consequently, assigned a NOB score of one. Mental rotation involve the PFC-coordinated activity in *both* the ventral and the dorsal visual cortices ^23,24,27^, and are therefore assigned the PCT score of two.

#### 5. Mental synthesis of several objects

##### 5.1 Spatial prepositions

Understanding of spatial prepositions such as in, on, under, over, beside, in front of, behind requires a subject to superimpose several objects. For example, the request “to put a green box {inside/behind/on top of} the blue box” requires an initial mental simulation of the scene, only after which is it possible to correctly arrange the physical objects. An inability to produce a novel mental image of the green box {inside/behind/on top of} the blue box would lead to the use of trial-and-error, which in majority of cases will result in an incorrect arrangement. Understanding an instruction to “put the bowl {*behind/in front of/on/under*} the cup requires the combination of two objects and therefore is assigned a NOB score of two. Understanding an instruction to “put the cup *in* the bowl and *on* the table” requires the combination of three objects and therefore is assigned a NOB score of three. In general, a NOB score in mental synthesis questions is defined as the number of disparate objects that have to be imagined together.

When two disparate objects are imagined together their spatial relationship has to be encoded. Encoding of spatial information is the function of the dorsal visual cortex ^23,24,27^. Accordingly, mental synthesis of multiple objects always involves the PFC-coordinated activity in *both* the ventral and the dorsal visual cortices and is therefore assigned the PCT score of two.

##### 5.2 Flexible syntax

The order of words, or syntax (from Greek *syn*, meaning *together*, and *taxis*, meaning *an ordering*), is an essential quality of all human languages. A change in the word-order often completely changes the meaning of a sentence. For example, the phrase “a cat ate a mouse” and the phrase “a mouse ate a cat,” have very different connotations only because the word order was changed. To understand the exact meaning of these two sentences, the subject must mentally synthesize the objects of “a cat” and “a mouse.”

Flexible syntax is distinguished from rigid syntax, whereby the meaning of a word is always dictated by its position in a sentence. For example, if we all agree that the first noun in the “noun1 ate noun2” sentence will always be the eater and the second noun will always be the prey, then the meaning of the sentence can be inferred without mental synthesis of multiple objects. When you hear “a cat ate a mouse,” you automatically know that noun1 (a cat) was the eater and noun2 (a mouse) was the prey. You do not need to synthesize the image of a cat and a mouse together; you can simply infer the meaning from the sequence of words in the sentence. Of course this rule constricts the conversation to a single sentence structure: “noun1 ate noun2.” If you follow this structure strictly, you will always misinterpret sentences like “a cat was eaten by a mouse,” “a cat will be eaten by a mouse,” “this cat never eats mice,” since you will always think that noun1 (the cat) is the eater. Human languages are not bound by limitations of rigid syntax: both the first and the second nouns can be the ‘eaters’ (“a cat ate a mouse” versus “a cat was eaten by a mouse”). All human languages use flexible syntax and rely on mental synthesis of multiple objects to decode the meaning of a sentence.

Similar to spatial prepositions tasks, a NOB score in flexible syntax tasks is defined as the number of disparate objects that have to be imagined together and PCT score of two is assigned.

##### 5.3 Before/after word order

To test understanding of before/after word order, subjects can be asked ‘to touch your nose after you touch your head.’ Similar to flexible syntax, this task rely on mental synthesis to simulate the process of touching the nose and the head. These tasks are given a NOB score of two and PCT score of two.

##### 5.4 Active vs. passive

Subjects can be asked to point to a picture showing ‘The boy is pulling the girl’ vs. ‘The girl is pulled by the boy.’ Similar to flexible syntax, this task relies on mental synthesis to combine the boy and the girl in the mind's eye. It is given a NOB score of two and PCT score of two.

##### 5.5 Exact arithmetic abilities

Mentally finding the solution to a complex arithmetic problem such as mental two-digit addition (e.g. 43+56) and multiplication (e.g. 32×24) also relies on mental synthesis. This process is different from multiplication of single digit numbers (e.g. 2×4) that can be retrieving from memorized multiplication table^28^. Since the exact number of objects in two-digit multiplication is not easily assertained, this task received an NOB score of “2+.” Mental two-digit addition and multiplication was assigned the PCT score of two.

#### 6. Special cases

##### 6.1. Classification tasks

Classification tasks involve sorting objects on the principles of gender, animacy, etc. Classification tasks involve no object combination and therefore were assigned a NOB score of one. The tasks also involve no modification of objects and therefore were assigned the PCT score of zero. Classification may be more challenging than similarly scored “Find the object” tasks, since the object's class has to be determined, but this is not reflected in the scoring system used in this report.

##### 6.2 Picture arrangement into a story

A typical test consists of three to six pictures with cartoon characters that can be arranged into a story. For example, picture A shows a boy running, picture B shows the boy falling, and picture C shows the boy crying. The pictures are presented to the subject in scrambled order with instructions to rearrange the pictures to make the best sense. A typical adult might approach this task by mentally animating the characters and generating a story using mental synthesis of multiple objects. However, in order to identify the minimal neurobiological requirements for the correct answer, we considered the following alternative. Most picture arrangement tasks rely on subject's understanding of causality between events shown in the pictures. The inference of causality can be made from a memory of similar events watched on TV or a personal experience, e.g. running, falling and crying thereafter. The process of inferring the causality does not rely on combination or modification of objects and therefore was assigned a NOB score of one and the PCT score of zero.

Comments:
1. The methodology for deriving a NOB and PCT scores from the performance IQ score was described in Ref ^1^.
2. Genie was tested with Raven's Matrices and Wechsler Intelligence Scale for Children (WISC). Fromkin *et al.* report that 1.5 years into rehabilitation Genie's “performance on the Raven Matrices could not be scored in the usual manner but corresponded to the 50th percentile of children aged 8.5 to 9 years” ^29^. Curtiss *et al.* report that 2 years into rehabilitation “on WISC… performance subtests … she has achieved subtest scaled scores as high as eight and nine” ^30^. In addition, Curtiss reports that 5 years into rehabilitation “Genie was given the Coloured Progressive Matrices Test. … Genie's overall score was 29.” ^6^ Using this limited information we attempted to map Genie's IQ into a NOB and PCT scores:
A. WISC uses three sections to determine the performance score: Matrix Reasoning, Figure Weights, and Visual Puzzles. A person who can only answer test questions with a NOB score of one and PCT score of zero (i.e. “Find the same object” and “Amodal completion”) would receive scaled section scores of 2, 5, 3 respectively ^1^. A person who can also answer test questions with a NOB score of one and PCT score of one (i.e. “Integration of size/color modifier”) would receive scaled section scores of 4, 5, and 4, respectively. A person who can also answer test questions with a NOB score of one and PCT score of two (i.e. “Integration of number modifier” and “Mental rotation and modification of object's location in space”) would receive scaled section scores of 6, 10, and 4, respectively. To demonstrate ability for mental synthesis of multiple objects, the subject has to have section scores above 6, 10, and 4, respectively. From the Curtiss *et al.* report that Genie” achieved subtest scaled scores as high as eight and nine” ^30^ we have to assume that the highest scaled section score was “eight and nine”. The easiest WISC section where Genie would be expected to show the highest score is the Figure Weights sections. The score of “eight and nine” in the Figure Weights section corresponds to a NOB score of one and PCT score of two. That is Genie demonstrated ability to “Find the same object,” “Integrate of size/color modifier,” “Integrate of number modifier,” and “Mentally rotate and modify object's location in space,” but failed questions that required mental synthesis of multiple objects.
B. From Fromkin *et al.* report that Genie's “performance on the Raven Matrices … corresponded to the 50th percentile of children aged 8.5 to 9 years” ^29^, assuming that Raven Standard Progressive Matrices test was administered and consulting the Raven manual ^31^, Table SPM12, we gather that Genie's raw score was between 33 and 36. This raw score corresponds to a NOB score of one and PCT score of two ^1^ and confirms that Genie demonstrated ability to “Find the same object,” “Integrate of size/color modifier,” “Integrate of number modifier,” and “Mentally rotate and modify object's location in space,” but failed questions that required mental synthesis of multiple objects.
C. The latest nonverbal test was administered 5 years into rehabilitation “Genie was given the Coloured Progressive Matrices Test … Genie's overall score was 29.” ^6^ Raven's Colored Progressive Matrices was originally designed for children aged 5 through 11 years-of-age, the elderly, and mentally and physically impaired individuals. This test contains sets A and B from the standard matrices, with a further set of 12 items inserted between the two, as set Ab. Most items are presented on a colored background to make the test visually stimulating for participants. The score of 29 means that Genie correctly answered 29 out of 36 questions. Assuming Genie has answered correctly the easiest question, we infer that she answered correctly all 12 questions in the set A (maximum NOB score of one and PCT score of zero), 6 questions in the set B (maximum NOB score of one and PCT score of one), and 11 questions in the set Ab (maximum NOB score of one and PCT score of zero) ^1^. In the Coloured Progressive Matrices Test the hardest questions Genie was able to answer had a NOB score of one and the PCT score of one. As Genie was not able to answer any questions with a NOB score of two, we conclude that Genie failed those questions that required mental synthesis of multiple objects. Thus, in all three nonverbal tests that Genie was administered up to five years into rehabilitation Genie has invariably failed to demonstrate capacity for mental synthesis of multiple objects.
3. Grimshaw reports that E.M.'s “performance on both simple and complex modification tasks was above chance but imperfect from the earliest assessment. E.M. made more errors in complex modification (two adjectives) than in simple modification (one adjective). Almost all errors involved a failure to identify the correct shape—the modifiers were always correct” ^32^.
4. Curtiss reports that “She didn't perform as well as Genie on such tasks, but she could put picture sequences into a logical sequence” (personal communications).
5. Curtiss reports that “She did not appear to comprehend (or use) such elements” (personal communications).
6. Curtiss reports that “On standardized (nonverbal) psychological and neuropsychological tests Chelsea consistently demonstrates a performance IQ of between 77 and 89” ^33^. It is unclear what exact nonverbal IQ tests were used with Chelsea. For most nonverbal tests, the score of 78 corresponds to a NOB score of one and PCT score of one; the score of 86 corresponds to a NOB score of one and PCT score of two ^1^. We conclude that at best Chelsea has demonstrated an ability to “Integrate number modifier,” and “Mentally rotate and modify object's location in space,” but failed those questions that required mental synthesis of multiple objects.
7. Morford reports that “Maria was administered Raven's Standard Progressive Matrices after three months and after 32 months exposure to (American Sign Language) ASL. Using norms gathered in 1986 for children in the US, her score after three months’ exposure to ASL was still below the fifth percentile for her age group, but after 32 months’ exposure to ASL, her performance was exactly at the fiftieth percentile for her age group” ^34^. Fifth percentile corresponds to the raw score of 35 and IQ score of 75 ^31^, a NOB score of one and PCT score of two ^1^. Fiftieth percentile corresponds to the raw score of 52 and IQ score of 100 ^31^, a NOB score of two and PCT score of two ^1^, indicating that Maria has correctly answered nearly 20 questions that required mental synthesis of multiple objects.
8. Morford reports that “Marcus was administered four of the WISC-R Performance Subtests after 19 and after 31 months exposure to ASL… Marcus's performance improved by more than ten scale points from the first to the second administration, increasing the corresponding performance IQ from 70.5 to 85 points. The WISC score of 70.5 corresponds to a NOB score of one and PCT score of zero ^1^. The WISC score of 85 corresponds to a NOB score of one and PCT score of between one and two. We conclude that at best Marcus has demonstrated and ability to “Integrate of number modifier,” and “Mentally rotate and modify object's location in space,” but failed in questions that required mental synthesis of multiple objects.
9. Subject I.C. was unable to use spatial signs (left/right, on top of/underneath) to describe the relationship between the objects.
10. According to Ramirez *et al.* ^35^, despite their age, the utterances of Shawna, Cody, and Carlos were neither long nor complex, never used lexical items indicating subordination or conditionals, and never used inflected verbs.
11. According to Ramirez *et al.* Cody received the score of 91 on the Test of Nonverbal Intelligence-3 (TONI-3) and the score of 85 on the Wechsler Nonverbal Scale of Ability ^36^ indicating that he failed to answer most or all questions that required mental synthesis of multiple objects.
12. According to Ramirez *et al.* Carlos received the score of 85 on the TONI-3 and the score of 74 on the Wechsler Nonverbal Scale of Ability ^36^ indicating that he failed to answer all questions that required mental synthesis of multiple objects.

## Results

Table 2 summarizes findings obtained from ten linguistic isolates. Genie (shown in first column) was linguistically isolated from 20 months of age until 13 years 7 months ^6,7^. After Genie was rescued, she was the subject of targeted attempts of language instruction. Genie became the subject of an investigation (funded by NIH) to discover if there was a critical age threshold for *first-language* acquisition. Within a few months she had advanced to one-word answers and had learned to dress herself. Her doctors predicted complete success. Her teachers included psychologists James Kent, linguist Susan Curtiss, therapist David Rigler, and many other well-intentioned people genuinely interested in rehabilitating Genie. After seven years of rehabilitation, Genie's vocabulary had expanded to several hundred words; however, she could only speak in short sentences with little use of syntax or verb tense. Genie's comprehension of complex sentence structures remained inconsistent. Remarkably, she was unable to grasp pronouns and prepositions such as *behind*, *in front*, *on* and *under* ^6,37^. Genie did not understand the difference between the simple task of “putting a bowl *behind* a cup” and “putting a bowl *in front of* a cup,” despite familiarity with both objects described by the words “bowl” and “cup.” These tasks, which are reasonably simple for a typical individual well before the age of six ^38^, were beyond Genie's capability even at the age of 20 after seven years of strenuous linguistic training. Thus, on linguistic tests Genie passed most tests with a NOB score of one, but did not pass any tests with a NOB score of two.

**Table 1.** The hierarchical classification of linguistic tests question by a NOB score and the PCT score.

| Linguistic test | Number of objects score | Posterior cortex territory score |
| --- | --- | --- |
| Find the object | 1 | 0 |
| Classification tasks | 1 | 0 |
| Picture arrangement into a story | 1 | 0 |
| Integration of size and color modifiers | 1 | 1 |
| Comparative: e.g. ‘Which button is smaller?’ | 1 | 1 |
| Superlative: e.g. ‘Which button is smallest out of five?’ | 1 | 1 |
| Integration of number modifier | 1 | 2 |
| Singular vs. plural | 1 | 2 |
| Understanding of simple negation | 1 | 2 |
| Mental rotation and modification of a single object’s location in space | 1 | 2 |
| Copying stick and block structures | 1 | 2 |
| Mental synthesis of multiple objects:<br>Understanding of spatial prepositions, such as in, on, under, over, beside, in front of, behind, etc. | 2 | 2 |
| Understanding of flexible syntax | 2 | 2 |
| Before/after word order: | 2 | 2 |
| Active vs. passive | 2 | 2 |
| Exact arithmetic abilities: mental two-digit addition / multiplication | 2+ | 2 |

**Table 2.** Linguistic isolates performance in verbal and nonverbal tests.

| NOB score | PCT score | TESTS | SUBJECTS | Genie | Victor of Aveyron | E.M. | Chelsea | Maria | Marcus | I.C. | Shawna | Cody | Carlos |
| --- | --- | --- | --- | --- | --- | --- | --- | --- | --- | --- | --- | --- | --- |
| 1 | 0 | Find the named object |  | Yes | Yes | Yes | Yes | Yes | Yes | Yes | Yes | Yes | Yes |
| 1 | 0 | Classification and categorization tasks |  | Yes | No data | No data | No data | No data | No data | No data | No data | No data | No data |
| 1 | 0 | Picture arrangement into a story |  | Yes | No data | Deficit | Yes<sup>4</sup> | No data | No data | No data | No data | No data | No data |
| 1 | 1 | Integration of size and color modifiers |  | Yes | No data | Yes<sup>3</sup> | No data | No data | No data | No data | No data | No data | No data |
| 1 | 1 | Comparative: e.g. 'Which button is smaller?' |  | Yes | No data | Yes | No data | No data | No data | No data | No data | No data | No data |
| 1 | 1 | Superlative: e.g. 'Which button is smallest out of five?' |  | Inconsistent | No data | Yes | No data | No data | No data | No data | No data | No data | No data |
| 1 | 2 | Negative vs. affirmative statements |  | Yes | No data | No | No data | No data | No data | No data | No data | No data | No data |
| 1 | 2 | Singular vs. plural |  | Yes | No data | Yes | No data | No data | No data | No data | No data | No data | No data |
| 1 | 2 | Understanding of simple negation |  | Yes | No data | No | No data | No data | No data | No data | No data | No data | No data |
| 1 | 2 | Copying stick and block structures |  | Yes | No data | No data | Yes | No data | No data | No data | No data | No data | No data |
| 2 | 2 | Understanding of spatial prepositions such as in, on, under, over, beside, in front of, behind, etc. |  | No | No data | No | No<sup>5</sup> | No data | No data | No<sup>9</sup> | No data | No data | No data |
| 2 | 2 | Understanding of flexible syntax |  | No | No | No | No | No | No | No data | No<sup>10</sup> | No<sup>10</sup> | No<sup>10</sup> |
| 2 | 2 | Before/after word order: e.g. 'Touch your nose after you touch your head.' |  | No | No data | No | No data | No data | No data | No data | No data | No data | No data |
| 2 | 2 | Active vs. passive: e.g. 'The boy is pulling the girl' VERSUS 'The girl is pulled by the boy' |  | No | No data | No data | No data | No data | No data | No data | No data | No data | No data |
| 2 | 2 | Exact arithmetic abilities: mental two-digit addition / multiplication |  | No data | No data | No data | No data | No data | No data | No | No data | No data | No data |
|  |  | Nonverbal (performance) IQ score and the derived NOB and PCT scores<sup>1</sup> |  | IQ score<sup>2</sup><br>NOB=1<br>PCB=2 | No data | IQ=83 to 88<br>NOB=1<br>PCB=2 | IQ=77 to 89<sup>6</sup><br>NOB=1<br>PCB=2 | IQ=73 to 100<sup>7</sup><br>NOB=1 to 2<br>PCB=2 | IQ=70.5 to 85<sup>8</sup><br>NOB=1<br>PCB=2 | IQ=65<br>NOB=1<br>PCB=0 | IQ=67<br>NOB=1<br>PCB=0 | IQ=84 to 91<sup>11</sup><br>NOB=1<br>PCB=2 | IQ=74 to 85<sup>12</sup><br>NOB=1<br>PCB=2 |

Genie was also tested on standardized nonverbal tests Raven's Matrices ^29,39^ and WISC ^30^. On both nonverbal tests Genie was able to answer questions with a NOB score of one but failed in questions with a NOB score of two (see comment 2 in the Table 2 legend). Thus, on both verbal and nonverbal tests Genie demonstrated ability to modify a single object but failed in both verbal and nonverbal questions that required mental synthesis of multiple objects.

Victor of Aveyron, an eighteenth-century French feral child entered his rehabilitation at the approximate age of 13 (Table 2, column 2). Jean Marc Gaspard Itard worked with the boy for five years teaching him words and recording his progress ^8^. Victor showed significant early progress in understanding and reading simple words, but failed to progress beyond a rudimentary level. Itard wrote, “Under these circumstances his ear was not an organ for the appreciation of sounds, their articulations and their combinations; it was nothing but a simple means of self-preservation which warned of the approach of a dangerous animal or the fall of wild fruit” ^8^. In a clear parallel to Genie's case, Victor of Aveyron learned many words after several years of training, but failed to acquire syntactic language.

Studies of feral children, however, are highly problematic: isolation and childhood abuse can result in all sorts of psychological problems, which may confound conclusions drawn about linguistic abilities. Studies of congenitally deaf people, who had linguistic but not social deprivation, have fewer methodological weaknesses. To date, there have been only eight carefully documented cases of deaf subjects who were linguistically deprived until puberty.

Gina Grimshaw and her colleagues studied a 19-year old man referred to as E.M., who has been profoundly deaf since birth and grew up in a rural area where he received no formal education and had no contact with the deaf community (Table 2, column 3). Although E.M. had not experienced formal language, he was exposed to simple communication. He and his family used homesign, a system of gestures that allowed them to communicate simple commands, but lacked much in the way of syntax ^32^. This is quite typical of families with deaf children and hearing parents, which are isolated from a sign language community; instead of learning a formal sign language, they normally spontaneously develop a home sign system. At the age of 15, E.M. was fitted with hearing aids that corrected his hearing loss, and he began to learn verbal Spanish. Grimshaw's tests demonstrated that E.M. had linguistic problems similar to those of Genie, Victor and other individuals who were linguistically deprived in their childhood.

While E.M.'s performance on simple linguistic tests that do not rely on mental synthesis was reasonably good, his performance on more complex tests that rely on mental synthesis of multiple objects was poor. Similar to Genie, E.M. had significant difficulty with spatial prepositions, even though his performance on this task was better than chance. Grimshaw *et al.* report that “even at the 34-month assessment, he had not mastered one of these prepositions, nor were his errors limited to related pairs (*under* vs. *over*, *in front of* vs. *behind*). His general strategy when following a direction to “put the green box *in* the blue box,” was to pick up the appropriate two boxes and to “move them through a variety of spatial arrangements, watching the examiner for clues as to which was correct” ^32^. Of course, normal performance of such a task would first involve a mental simulation of the green box *inside* the blue box in the process of mental synthesis, followed by a demonstration of the results of the mental simulation with the physical objects. The nonverbal tests also indicate that E.M. was not able to answer questions with a NOB score of two. At age 16, E.M's IQ was estimated to be 83 with the Test of Nonverbal Intelligence (TONI-II) and 88 with the Performance Scale of the Wechsler Intelligence Scale for Children-Revised (WISC-R). Reassessment with the Wechsler Adult Intelligence Scale—Revised (WAIS-R) at age 19 produced a Performance IQ of 85. Thus, on both verbal and nonverbal tests E.M. could not demonstrate an ability for mental synthesis of multiple objects.

Another report was of Chelsea, a deaf woman isolated from syntactic linguistic input until she was 32, when her first exposure to spoken English was provided by successful auditory amplification (Table 2, column 4). Similar to E.M., Chelsea used homesign to communicate until the auditory amplification apparatus was installed ^39^. With adequate auditory amplification, Chelsea managed to learn many words. However, Chelsea never learned to put those words together into syntactically meaningful phrases, and she could not understand syntactically complex sentences. Similar to Genie, Chelsea did not appear to comprehend or use spatial preposition. She also did not seem to understand directions that relied on spatial prepositions such as “put a bowl *behind* a cup” (NOB score of two, Curtiss S, personal communications). On nonverbal IQ tests Chelsea scored between 77 and 89 ^33^ indicating that she failed questions with a NOB score of two. Thus, on both verbal and nonverbal tests Chelsea could not demonstrate an ability for mental synthesis of multiple objects.

Jill Morford studied two deaf adolescents, Maria and Marcus, as they were acquiring their first language (American Sign Language, or ASL), at the ages of 13 and a half and 12 respectively (Table 2, columns 5 and 6). Both Maria and Marcus were born profoundly deaf ^34^. They were not raised in isolation and already had well developed social and cognitive skills at the time of their first exposure to ASL. Their first exposure to ASL happened after their families emigrated from countries with limited educational resources for deaf children to North America where public education for deaf children is mandatory. After approximately seven years of exposure to ASL, Maria and Marcus were tested by Jill Morford. The comprehension of spatial prepositions was not studied (Morford J, personal communications), but the examination included another test for understanding of flexible syntax (NOB score of two). Maria and Marcus were shown eight sentences describing events of climbing, falling, looking, biting, and licking. To show their understanding of a sentence, Maria and Marcus had to select a correct picture from a set of four pictures. The correct picture depicted the event that the target sentence described, and the three incorrect pictures depicted events that were related in path movement and/or protagonists. For example, “for the target sentence describing the boy climbing a tree, distracter pictures depicted the boy climbing a rock surrounded by trees, the frog climbing out of a jar, and the boy and the dog looking over a fallen tree” ^34^. In order to select the correct response, the subjects had to mentally synthesize the scene described by the sentence and then select the picture best reflecting the result of their mental synthesis. Chance performance for the comprehension task is 25%. Both participants selected the correct picture 38% of the time, barely above chance level. Even when the participants were allowed to review the topic sentence an unlimited number of times, Maria still selected the correct picture at the rate of only 63%.

Maria was administered Raven's Standard Progressive Matrices test after three months and after 32 months exposure to ASL. Her score after three months’ exposure to ASL was below the fifth percentile for her age group, corresponding to a NOB score of one and the PCB score of two. After 32 months’ exposure to ASL, her performance increased to the fiftieth percentile for her age group ^34^, corresponding to a NOB score of two and PCB score of two. Marcus was administered four of the WISC-R Performance Subtests after 19 and after 31 months exposure to ASL, including picture completion, picture arrangement, block design, and object assembly. Marcus's performance IQ score (computed by prorating from these four tests) was 70.5 and 85 points respectively ^34^, indicating that he failed questions with a NOB score of two.

Daniel C. Hyde and colleagues studied the cognitive abilities of a 13-year-old deaf child named I.C. (Table 2, column 7). As a result of living in an underdeveloped country, I.C. received no formal sign language instruction until the age of 13 when he immigrated permanently to the United States ^40^. I.C. communicated his thoughts and emotions mostly through facial expression and by a homesign gestural system developed inside the family. Following his enrollment in the residential program for deaf children, he quickly learned a great number of ASL signs. The experiments were performed using both the ASL and the homesign system (the homesign system was used with the help of I.C.'s younger brother translating the tasks to I.C.). I.C. was presented with pictures containing pairs of objects: side by side or one on top of the other and was asked to describe the pictures. I.C. had no trouble naming the objects presented to him, but was unable to use spatial signs (left/right, on top of/underneath) to describe the relationship between the objects. Even after the experimenter prompted I.C. with a correct response following each attempt (for example, the experimenter signed “dog on top of cat” or “dog on the left/ cat on the right”), I.C. was still unable (in the following attempt) to “describe a picture in a manner that allowed the spatial relationship to be unambiguously interpreted” ^40^. The researchers concluded that “tests of spatial and geometrical abilities revealed an interesting patchwork of age-typical strengths and localized deficits. In particular, the child performed extremely well on navigation tasks involving geometrical or landmark information presented in isolation, but very poorly on otherwise similar tasks that required the combination of the two types of spatial information.” I.C. was tested with Wechsler Intelligence Scale for Children-IV (WISC-IV). His nonverbal IQ score was 65, indicating that he failed questions with a NOB score of two. Thus, on both verbal and nonverbal tests I.C. failed to demonstrate an ability for mental synthesis of multiple objects.

Naja Ferjan Ramírez and colleagues studied the acquisition of American Sign Language in three adolescents (Shawna, Cody and Carlos) who, similarly to other deaf linguistic isolates, were deprived of a formal sign language until the age of 14 (Table 2, columns 7 to 9). Prior to receiving training in ASL, these subjects, who grew up in separate families, relied on behavior and a limited number of gestures to communicate ^35^. The researchers did not conduct any linguistic tests that could be used to judge the subjects’ mental synthesis. However, subjects’ utterances used in spontaneous conversations are consistent with significantly diminished mental synthesis. Despite their age, their utterances were neither long nor complex, never used lexical items indicating subordination or conditionals, and never used inflected verbs ^35^. Subjects “used few if any spatial prepositions or ASL classifiers describing spatial relations” (Mayberry, personal communications). Their scores on the TONI-3 were 67 (Shawna), 91 (Cody) and 85 (Carlos). Cody and Carlos were also tested on the Wechsler Nonverbal Scale of Ability ^36^. Cody and Carlos scored 85 and 74, respectively. These nonverbal IQ scores indicate that subjects failed questions with a NOB score of two and confirm subjects’ difficulties with mental synthesis of multiple objects.

## Discussion

In this manuscript we analyzed all the published reports of linguistic isolates – individuals not exposed to syntactic language until puberty. The stark contrast between their lifelong difficulty in understanding spatial prepositions and syntax and their ability to learn hundreds of new words begs the question: is lack of syntactic language comprehension a language problem or an imagination problem? If linguistic isolates were not able to purposefully synthesize disparate objects into novel configurations in their mind's eye, that could parsimoniously explain their lack of understanding flexible syntax, low nonverbal IQ, and their lifelong difficulty of processing spatial information. To test this hypothesis, we noticed that mental synthesis of disparate objects is the function of the lateral prefrontal cortex (LPFC) ability to control the posterior cortex neurons ^1^ and ranked all verbal and nonverbal tests performed by linguistic isolates by: a) the number of disparate objects that have to be combined together by the LPFC in order to answer the question and b) the amount of physical territory within the posterior cortex that needs to be recruited by the LPFC. According to our analysis, linguistic isolates performed well in all tests that did not involve the LPFC control of the posterior cortex, showed decreasing scores in tests that involved greater recruitment of the posterior cortex by the LPFC, and failed in tests that involved the greatest recruitment of posterior cortex necessary for mental synthesis of multiple objects. This performance pattern is in line with observations in late *first-language* learners ^14–16,41^, individuals with nonverbal ASD ^42^, as well as in patients with LPFC damage who are often unable to correctly answer questions that require integration of modifiers, mental rotation and modification of an object's location in space, as well as mental synthesis of several objects ^43–47^.

All of these observations are consistent with the hypothesis that understanding complex syntactic language is intimately connected to the control of the LPFC over objects encoded in the brain.

### Objects as units of perception

Our visual world consists of meaningful, unified, and stable objects that move coherently as one piece. Objects, therefore, constitute the functional units of perception ^48^. Over time, neurons synchronously activated by visual observation of an object, get wired together by enhanced synapses and form a resonant system that tends to activate as a single unit resulting in perception of that object. These resonant networks of neurons encoding objects in the cerebral cortex are known as neuronal ensembles ^49^. When one perceives any object, the neurons of that object's neuronal ensemble activate into synchronous resonant activity ^50^. The neuronal ensemble binding mechanism, based on the Hebbian principle “neurons that fire together, wire together,” came to be known as the binding-by-synchrony hypothesis ^51,52^.

While the term “neuronal ensemble” is often used in a broad sense to refer to any population of neurons involved in a particular neural computation, in this discussion we will use the term more narrowly to mean a stable group of neurons which are connected by enhanced synapses and which encode specific objects. Thus, neuronal ensembles are internal equivalents of objects: while the latter are physical or external units of perception, the former are the internal units of perception; each object that has been seen and remembered by a subject is encoded by a neuronal ensemble.

The neurons of an ensemble are distributed throughout the posterior cortex (occipital, temporal, and parietal lobes). Most neurons encoding a stationary object are located within the ventral visual cortex (also known as the "what” stream) ^21^, Fig. 1. These thousands of neurons encode the various characteristics of each object, such as shape, color, texture, etc. ^53,54^. The majority of neurons encoding one's favorite cup are located in the primary visual area (V1). A smaller number of more specific neurons are located in the extrastriate areas such as V2 and V4, and an even smaller number of the most specific neurons are located in the temporal lobe. Most humans can use their PFC to purposefully change elements of a neuronal ensemble enabling an alteration of an object's color, size, rotation, or position in space within the “mind's eye.”

### The role of the PFC

The PFC consists of two functionally distinct divisions. The ventromedial PFC is predominantly involved in emotional and social functions such as the control of impulse, mood, and empathy ^55^. The lateral PFC (LPFC) is predominantly, but not exclusively, involved in “time integrating and organizing functions, such as control of working memory and imagination” ^56^. The LPFC is evolutionarily related to the motor cortex and there are a number of pertinent parallels between the two ^55,56^. Just as the primary motor cortex can recruit motor units of voluntary muscles, the LPFC is able to activate neuronal ensembles in the posterior cortex. If motor units are homologous to neuronal ensembles and the LPFC is homologous to the motor cortex, then movement of muscles is homologous to “movement” of thoughts. The underlying mechanisms behind this “movement” of thoughts and their relationship to syntactic language are the primary focus of the following discussion.

### Lateral PFC-driven synchronization of neurons as the likely mechanism of internally-driven novel sensory experiences

We have previously hypothesized that the mechanism behind the mental synthesis of independent objects involves the LPFC acting in the temporal domain to *synchronize* the neuronal ensembles which encode those objects ^57,58^. Once these neuronal ensembles are time-shifted by the LPFC to fire *in-phase* with one another, they are consciously experienced as a unified object or scene and could therefore be mentally examined as a cohesive unit. This hypothesis is known as Mental Synthesis theory ^57^ and it is indirectly supported by several lines of experimental evidence ^59–63^.

We have also hypothesized that the neurological process of modifier integration, whereby the LPFC modifies the activity of neurons in a single neuronal ensemble resulting in changes of an object's perceived color or size, involves the LPFC acting in the temporal domain to *synchronize* the color encoding neurons in the visual area V4 with the object's neuronal ensemble in the ventral visual cortex ^1^. Once these neurons in V4 are time-shifted by the LPFC to fire *in-phase* with the object's neuronal ensemble, the object is consciously experienced in the new color. LPFC-driven synchronization of neurons in the posterior cortex is likely a general mechanism necessary to achieve any *internally-driven novel* sensory experience. The LPFC can be viewed as a puppeteer and the objects encoded in the posterior cortex as its many puppets. By pulling the strings, the LPFC changes the firing phase of the retrieved neuronal ensembles, synchronizing them into new mental constructs. Thus, understanding of syntactic language requires the LPFC puppeteer to precisely schedule the firing phase of the neuronal ensembles puppets dispersed throughout the posterior cortex.

### Synchronous frontoposterior connections

The Mental Synthesis theory also predicts that to synchronize neuronal ensembles dispersed throughout the posterior cortex, the LPFC must rely on synchronous connections to posterior cortex neurons ^1,58^. In many animal models synchronous connections have been observed between multiple brain areas that depend on precise timing for communication despite varying path lengths. In the cat retina, axons from peripheral regions have a greater conduction velocity than axons from neurons at the center of the retina to assure the simultaneous arrival of impulses in the brain ^64^. Experiments in rats show that myelination is the primary factor producing uniform conduction time (to within 1ms) in connections between the inferior olive nucleus in the brainstem and the cerebellar cortex, despite a wide variation in axon length ^65,66^. In cats, synchronous activation of groups of cells distributed in distant cortical locations has been shown in the visual cortex ^67^ and even between the two hemispheres of the brain ^68^. It was also shown in rats that synchronous action potential delivery from the layer V pyramidal neurons in the ventral temporal lobe to target subcortical regions is based on the differential conduction velocity in each fiber ^69^, which appears to be best explained by differential myelination changing the conduction velocity of the individual axons ^70^. The synchronous connections achieved by differential myelination were also observed in cats between the amygdala and the perirhinal cortex ^71^ and in mice between the thalamus and the somatosensory cortex ^72^. Accordingly, we hypothesized that synchronous frontoposterior connections achieved by differential myelination are essential for the LPFC's ability to synchronize posterior cortex neurons located at various physical distances from the LPFC and that these connections are therefore essential for understanding of complex syntactic language, Fig. 3 ^58^.

**Figure 3.**
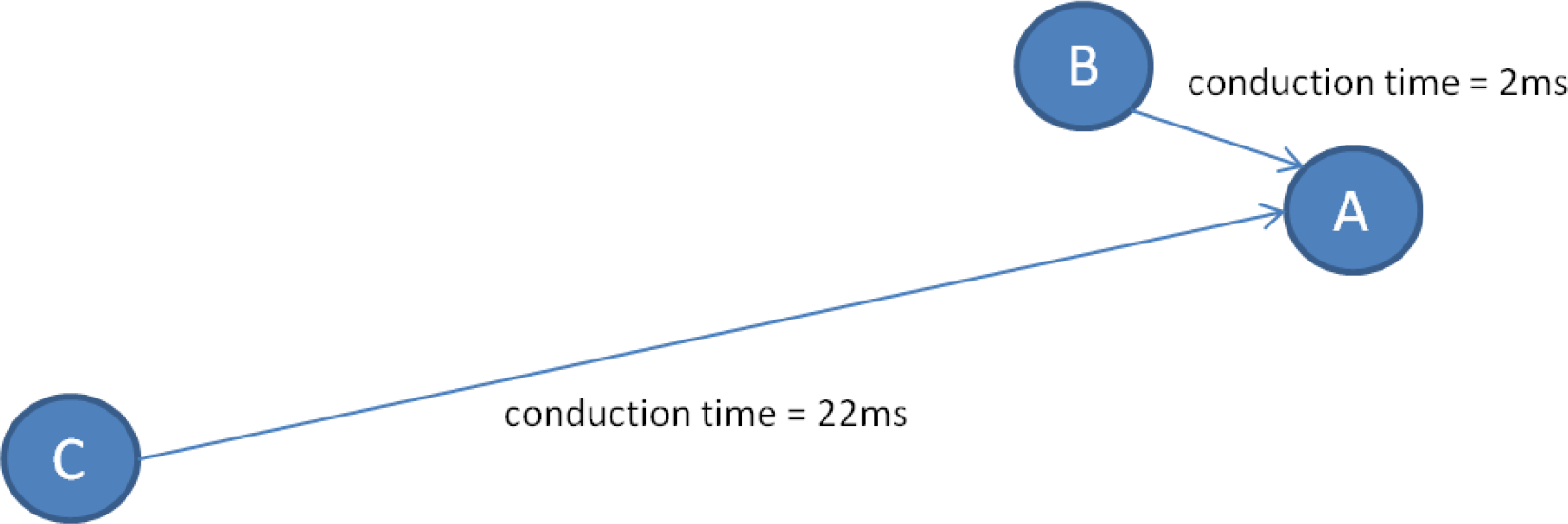
*Synchronicity* has to be understood in terms of synchronicity of the arrival of action potentials to a target neuron rather than absolute equality of action potential conduction times over different paths. Consider the following example: suppose neuron A is receiving excitatory input from neurons B and C via two different pathways (neuron A is the target neuron for both neurons B and C). Suppose that the action potential conduction time is 2ms from neuron B to neuron A and 22ms from neuron C to neuron A (i.e., the axonal pathway B-A has a significantly shorter conduction time than the axonal pathway C-A). Does it mean that the connections B-A and C-A are always asynchronous? No. The answer depends on the predominant neural activity rhythm in this network. At the firing rate of 50Hz (inter-spike interval of 20ms that correspond to Gamma rhythm), neurons B and C can actually be considered synchronous in relationship to neuron A: consider a train of action potentials synchronously fired by neurons B and C. The first action potential from neuron B will reach neuron A in 2ms and the first action potential from neuron C will reach neuron A in 22ms. Obviously, there would be no coincidence in the arrival times of the 1st action potentials from neurons B and C. However the second action potential from neuron B will arrive to neuron A in 22ms, concurrently with the 1st action potential from neuron C. Thus, starting with the second action potential, neuron A will receive synchronous activation from neurons B and C. The synchronous activation has a significantly greater probability of enhancing synaptic connections between neurons A and B, and A and C (Hebbian learning: ‘neurons that fire together, wire together’ ^49^). Thus, synchronicity does not need to imply absolute equality in the conduction time over different pathways. Rather synchronicity implies near-zero phase-shift between the two firing trains of action potentials at the postsynaptic cells. This phase-shift depends on conduction times over each pathway and also on the dominant firing frequency in the neural network.

### Ontogenetic acquisition of synchronous frontoposterior connections

Myelination is nearly completed by birth in most species in which the young are relatively mature and mobile from the moment of birth, such as wild mice ^73^ and horses ^74^. However, in humans, myelination is delayed considerably. Few fibers are myelinated at birth and some brain regions continue myelination well into mid-life. The PFC region is the last region to complete myelination. Myelination of cortico-cortical fibers that connect the PFC to other cortical areas does not reach full maturation until the third decade of life or later ^75,76^. To confer an evolutionary benefit, this delay in myelination must be associated with a practical advantage. Humans’ delayed maturation of myelination raises the possibility that experience determines myelination in a manner that optimizes synchronicity in neural networks.

There is significant evidence that neural activity can affect myelination. The number of myelin-forming cells in the visual cortex of rats increases by 30% when the rats are raised in an environment that is enriched by additional play objects and social interaction ^77^. Premature eye opening in neonatal rabbits increases myelin production presumably by increasing activity in the optic nerve ^78^. In contrast, rearing mice in the dark reduces the number of myelinated axons in the optic nerve ^79^. The proliferation of myelin-forming precursor cells in the rat's developing optic nerve depends on electrical activity in neighboring axons ^80^. In neuronal cell culture, induction of action potentials with electrical stimulation affects axonal myelination ^81,82^.

Environmental effects on axonal myelination extend beyond animal studies. In human infants, early experiences increase myelination in the frontal lobes in parallel with improved performance on cognitive tests ^83^. On the contrary, children suffering from abuse and neglect show on average a 17% reduction in myelination of the corpus callosum, the structure comprised of connections between the neurons of the left and the right hemispheres of the brain ^84^. Extensive piano practicing in childhood is accompanied by increased myelination of axons involved in musical performance, the white matter thickening increases proportionately to the duration of practice ^85^.

To sum up, synchronicity in at least some neuronal networks seems to be achieved via differential myelination and myelination may be experience-dependent. In fact, **considering the many variables affecting conduction delays in an adult brain and the hundreds of millions of frontoposterior connections, genetic instruction alone would seem inadequate to specify the optimal conduction velocity in every axon**. **Accordingly, the development of synchronous frontoposterior connections likely involves adjustment of the conduction velocity in individual fibers via an experience-dependent process.**

The experience-dependent developmental mechanisms have been clearly demonstrated for the visual, auditory, somatosensory and motor systems in virtually all species, from humans to Drosophila ^86,87^. Childhood sensory deprivation, such as monocular deprivation ^88–93^ or monaural plugging ^94^ and alterations in sensory input, such as caused by strabismus ^95^, rearing in a restricted auditory environment, or misalignment of the auditory and visual space maps ^96,97^, result in lifelong changes of neural networks and a deficit in the corresponding function. On the contrary, increased sensory experience have been associated with improved neural connections: in humans, musical training in childhood leads to an expanded auditory cortical representation and increased myelination of motor axons involved in musical performance ^85,98^ and introduction to a second language before the age of 12 results in a more optimal language encoding ^99,100^.

A common thread between these experience-dependent processes is that the ontogenetic experience essential for the formation of the neurological network is the same experience for which this network is naturally used in adulthood: stimulation of eyes by light is essential for formation of visual cortical connections, auditory stimulation is essential for formation of auditory cortex connections, motor practice is essential for formation of cortical motor connections, etc. Thus, it is compelling to hypothesize that cortical connections, which are essential for mental synthesis of multiple objects in adulthood are acquired as a result of the childhood mental synthesis exercises. Noting that conscious purposeful mental synthesis of disparate objects is primarily stimulated by syntactic language – whether it is internal language when we are thinking to ourselves or external language when we are involved in conversation – we would predict that the synchronous frontoposterior connections can only be acquired as a result of childhood exposure to syntactic language. The lack of syntactic language during the sensitive period of the PFC plasticity would result in asynchronous frontoposterior connections and mental synthesis disability.

In fact, this is exactly what we have observed in all linguistic isolates. Their lifelong mental synthesis disability is consistent with the critical role of early syntactic language use for establishment of synchronous frontoposterior connections. Childhood use of syntactic language (both externally and internally) likely provides the necessary training mechanism for development of synchronous connections between the LPFC and the sensory memory stored as neuronal ensembles in the posterior cortex. Without those synchronous connections having developed before the closure of the period of plasticity, the LPFC executive remains forever disabled at performing precise phase control of neurons, which are located over the large parts of the posterior cortex. The millions of strings connecting the LPFC puppeteer to its puppets (the neuronal ensembles in the posterior cortex) remain ill-adjusted, with the LPFC unable to control synchronization of independent neuronal ensembles and, therefore, unable to synthesize novel mental images. A child involved in a normal syntactic conversation (either with another person or with himself) internally manipulates mental images in his/her mind and during this process adjusts conduction velocity to achieve synchronicity in the neural networks connecting the LPFC to widespread regions of the posterior cortex.

The critical role of the syntactic language in formation of synchronous frontoposterior connections and mental synthesis is consistent with findings in late *first-language* learners and nonverbal individuals with autism. Elissa Newport and Rachel Mayberry studied the acquisition of sign language in deaf individuals differing in age of exposure: some were exposed to sign language from birth, while other children first learned sign language at school. These studies were conducted with adults who have been using sign language for at least twenty years. The results of the studies consistently indicated a negative correlation between the age of sign language acquisition and ultimate proficiency: those exposed to sign language from birth performed best, and late learners — worst, on all production and comprehension tests ^101,102^. Furthermore, late *first-language* learners have syntactic difficulty describing events with complex temporal relations ^103^, and have slower and less accurate performance in mental rotation independent of age and the total number of years of experience with the language ^14–16,41^. Emmorey *et al.* found that adults who had learned American Sign Language (ASL) early in life were more accurate than later learners at identifying whether two complex-shape figures presented at different degrees of rotation were identical or mirror images of each other ^41^. Martin *et al.* found that men who learned ASL natively were faster than later learners at identifying whether two-dimensional body-shaped figures (bears with one paw raised) presented at different rotations were identical or mirror images of each other ^14^. Martin *et al.* demonstrated that even after decades of signing experience, the age at which one first learned Nicaraguan Sign Language was a better predictor of mental rotation accuracy than the total number of years of experience with the language ^15^. Across all ages, early learners were more accurate than late learners. Pyers *et al.* studied generational differences between the 2 cohorts of deaf Nicaraguan signers who learned the emerging sign language at the same age but during different time periods – the first cohort of signers who grew up when the sign language was in its initial stage of development with few spatial prepositions and the second cohort of children who grew up a decade later exposed to more spatial prepositions and other recursive elements ^16^. The study used two spatial tasks that required the use of a landmark to find a hidden object: in the disoriented search condition, participants entered a small, enclosed room with a single red wall as a landmark. They watched an experimenter hiding a token in one corner, and then were blindfolded and turned slowly until disoriented. Then the blindfold was removed and the subjects indicated the corner where they thought the token was hidden. In the rotated box condition, the procedure was similar: the token was hidden in a small-scale tabletop model of the room, the participant was blindfolded, and the model (not the participant) was turned. The second-cohort signers, now in their 20s outperformed the first cohort, now in their 30s, both when they were disoriented and when the box was rotated. The first-cohort participants made their decisions slowly and with effort, taking as long as nine seconds and reporting that the task was difficult. Even in neurotypical children, production of spatial terms from 14–46 months predicted their performance on nonlinguistic spatial tasks at 54 months ^104^. Better and faster performance in spatial tasks in individuals exposed to more complex syntactic language earlier in life is best explained by the crucial role of syntactic language in fine-tuning the frontoposterior connections between the LPFC and the posterior cortex.

Prelingual deafness is such a serious concern that the US government has enacted laws to identify affected newborns. In 1999, the US congress passed the “Newborn and Infant Hearing Screening and Intervention Act,” which gives grants to help states create hearing screening programs for newborns. Otoacoustic Emissions Testing is usually done at birth, followed by an Auditory Brain Stem Response if the Otoacoustic Emissions test results indicated possible hearing loss. Such screening allows parents to expose deaf children to a formal sign language as early as possible and therefore avoid any delay in introduction to syntactic language.

Problems observed in late *first-language* learners, also afflict many individuals with autism, leading to what is commonly described as “stimulus overselectivity,” “tunnel vision,” or “lack of multi-cue responsivity” ^38,105–107^. Thirty to forty percent of individuals diagnosed with autism experience lifelong impairment in the ability to understand flexible syntax and spatial prepositions, and to mentally solve even the simplest of problems^108^. These individuals, commonly referred to having low-functioning ASD, typically exhibit full-scale IQ below 70 ^42,109^ and perform below the score of 85 in non-verbal IQ tests (see Ref. ^42^, Table 14.1, Performance Quotient). Since, significant language delay is a common problem in children with autism, lack of childhood use of syntactic language is likely one of the main contributing factors for this disability.

#### The duration of the critical period for acquisition of synchronous frontoposterior connections

The period of plasticity for the development of synchronous frontoposterior connections seems to close in several stages. Based on the randomized controlled study of institutionalized Romanian children, the plasticity seems to already decline before the age of two: children placed in foster care and therefore exposed to complex syntactic language before the age of 2 showed increased myelination and performed better in mental integration tasks compared to children who have been placed in foster care after the age of 2 ^18^.

The next significant decline in plasticity is around the age of five. When the left hemisphere is surgically removed before the age of five (to treat cancer or epilepsy), patients often attain normal cognitive functions in adulthood (using the one remaining hemisphere). Removal of the left hemisphere after the age of five often results in significant impairment of syntactic language and tasks requiring mental synthesis ^4,110–113^. The decline of plasticity after the age of 5 likely corresponds to a consolidation of asymmetry between hemispheres with one of the hemispheres (usually the left one) taking most of the control of the language function with underlying left LPFC puppeteer synchronously connected to the rest of the cortex.

Further reduction of plasticity occurs by the time of puberty; a lack of experience in syntactic speech before puberty likely results in a consolidation of widely desynchronized frontoposterior networks and prevents individuals from subsequent acquisition of mental synthesis. While parts of the PFC network retain some plasticity for a significantly longer period of time, since myelination of the PFC continues into the third decade of life or later ^75^, this plasticity seems inadequate to assist patients who were linguistically deprived until puberty in the acquisition of mental synthesis despite many years of intensive therapy.

### Is there a neurological dissociation between language and cognition?

It is common among linguists to discuss the “language module” in the brain. In support of the existence of the “language module,” linguists enlist patients with apparent dissociation between language and cognition ^37^. Observations in low functioning patients with fluid language production are pitted against the observations in high functioning individuals with aphasia. Is there real neurological dissociation between language and cognition and an easily separable language module? The answer to these questions depends on the definition of language.

The word “language” has highly divergent meanings in different contexts and disciplines. In informal usage, a language is usually understood as a culturally specific communication system (German, French, or Spanish Language), i.e. understanding of individual words without any reference to understanding of flexible syntax. On the other hand, in linguistics, the term “language” is used quite differently to encompass both understanding of words and flexible syntax. Fitch, Hauser, & Chomsky ^114^ explain the ambiguity of the “language” definition: “It rapidly became clear in the conversations leading up to HCF (i.e. Ref. ^115^) that considerable confusion has resulted from the use of “language” to mean different things. We realized that positions that seemed absurd and incomprehensible, and chasms that seemed unbridgeable, were rendered quite manageable once the misunderstandings were cleared up. For many linguists, “language” delineates an abstract core of computational operations, central to language and probably unique to humans. For many biologists and psychologists, “language” has much more general and various meanings, roughly captured by “the communication system used by human beings.” Neither of these explananda are more correct or proper, but statements about one of them may be completely inapplicable to the other.”

Mental synthesis mediated by the LPFC is the best candidate for “abstract core of computational operations, central to language and probably unique to humans” described by Fitch, Hauser, & Chomsky ^114^. Mental synthesis is an essential part of all human languages. When we speak we use mental synthesis to describe a novel image (“My house is the second one on the left, just across the road from the red gate”) and we rely on the listener to use mental synthesis in order to visualize the novel image. When we tell stories, we are often describing things that the listener has never seen before (“That creature has two heads, three tails, dark eyes, and can run faster than a cheetah”) and we rely on the listener to imagine the story in their mind's eye. As Steven Pinker put it, “the speaker has a thought, makes a sound, and counts on the listener to hear the sound and to recover that thought” ^116^.

All human languages are recursive: they allow *high fidelity transmission* of *infinite* number of *novel* images with the use of a *finite* number of words, Fig. 4. This is only possible because both listeners and speakers have mental synthesis ability presumably mediated by synchronous frontoposterior connections. Thus, both syntactic language and cognition depend on mental synthesis and on synchronous frontoposterior connections. The dissociation observed in many patients ^37^ is not between syntactic language and cognition, but rather between their use of words (expressive language) and their mental synthesis ability often assessed by measuring their nonverbal IQ ^1^. In fact, expressive language outside of a complex dialog cannot reliably serve as an indicator of mental synthesis ability. The following examples illustrate the problem of expressive language interpretation:

1. Some expressive language is completely automated, for example, coprolalia sometimes observed in patients with Tourette syndrome involves spontaneous and articulate uttering of expletives in an uncontrollable manner ^117^.
2. Tape recorder language and reading poetry by heart can sometimes be completely automated, prerecorded through repetition ^37,118^.
3. Echolalia in its profound form is automatic and effortless, imitative behavior whereby sounds or actions are imitated without explicit awareness ^119,120^.
4. Confabulations involve the production of fabricated memories without the conscious intention to deceive. A liar would commonly initially simulate a story by using LPFC-controlled mental synthesis of multiple objects and check it for consistency before delivering the story to the listeners. The confabulator, by definition, does not have a pre-simulated story that could indicate the ability of the LPFC to ascertain control over the posterior cortex. Confabulatory tendencies are often interpreted as an indicator of a disconnection between PFC and the posterior cortex ^121^.
5. Many people report the experience of reading aloud a book to a child and thinking of something else. In this case their LPFC is likely driving their mental synthesis while the book's sensory input is driving the Broca's area speech production and the right hemisphere's prosody in the bottom-up manner.
6. The vivid, original, bizarre, emotional, and story-like dreams that most people remember when they wake up occur during rapid eye movement (REM) sleep. These dreams consist of existing fragments of memory put into various new combinations and are often erroneously assumed to be created by the LPFC via the mechanism similar to mental synthesis. For example, Darwin associates imagination with dreaming: “The imagination is one of the highest prerogatives of man. By this faculty he unites former images and ideas, independently of the will, and thus creates brilliant and novel results. … Dreaming gives us the best notion of this power… The value of the products of our imagination depends of course on the number, accuracy, and clearness of our impressions, on our judgment and taste in selecting or rejecting the involuntary combinations, and to a certain extent on our power of voluntarily combining them. As dogs, cats, horses, and probably all the higher animals, even birds have vivid dreams, and this is shown by their movements and the sounds uttered, we must admit that they possess some power of imagination.” ^122^

**Figure 4.**
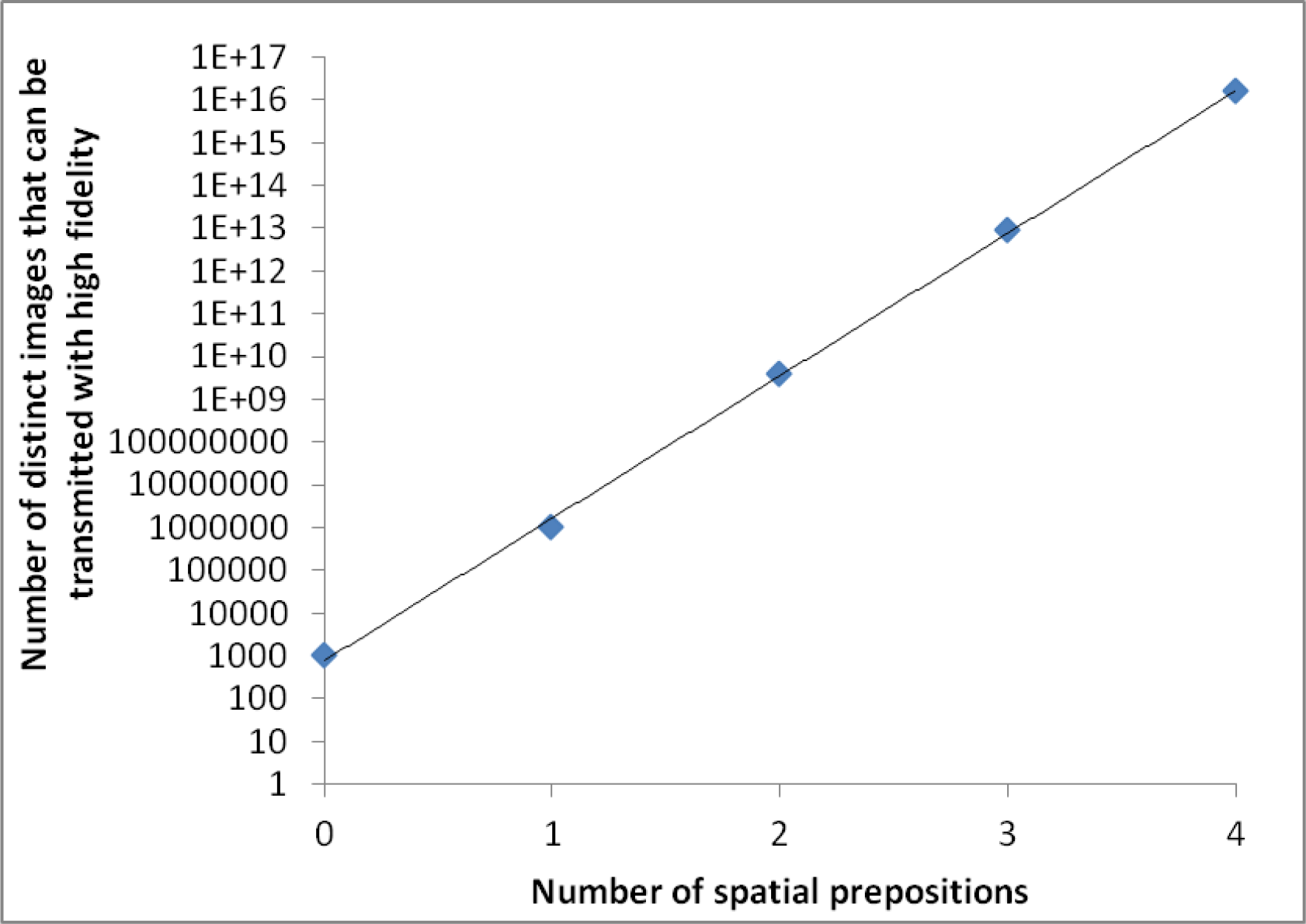
Flexible syntax, prepositions, adjectives, verb tenses, and other common elements of grammar, all facilitate the human ability to communicate an infinite number of novel images with the use of a finite number of words. The graph shows the number of distinct images that can be transmitted with high fidelity in a communication system with 1,000 nouns as a function of the number of spatial prepositions. In a communication system with no spatial prepositions and other recursive elements, 1000 nouns can communicate 1000 images to a listener. Adding just one spatial preposition allows for the formation of three-word phrases (such as: ‘a bowl *behind* a cup’ or ‘a cup *behind* a bowl’) and increases the number of distinct images that can be communicated to a listener from 1000 to one million (1000×1×1000). Adding a second spatial preposition and allowing for five-word sentences of the form *object-preposition-object-preposition-object* (such as: a bowl *on* a cup *behind* a plate) increases the number of distinct images that can be communicated to four billion (1000×2×1000×2×1000). The addition of a third spatial preposition increases the number of distinct images to 27 trillion (1000×3×1000×3×1000×3×1000), and so on. In general, the number of distinct images communicated by three-word sentences of the structure object-preposition-object equals the number of object-words times the number of prepositions times the number of object-words. A typical language with 1000 nouns and 100 spatial prepositions can theoretically communicate 1000^101^ × 100^100^ distinct images. This number is significantly greater than the total number of atoms in the universe. For all practical purposes, an infinite number of distinct images can be communicated by a syntactic communication system with just 1000 words and a few prepositions. Prepositions, adjectives, and verb tenses dramatically facilitate the capacity of a syntactic communication system with a finite number of words to communicate an infinite number of distinct images. Linguists refer to this property of human languages as *recursion*. The “infiniteness” of human language has been explicitly recognized by “Galileo, Descartes, and the 17th-century ‘philosophical grammarians’ and their successors, notably von Humboldt” ^115^. The infiniteness of all human languages stand in stark contrast to finite homesign communication systems that are lacking spatial prepositions, syntax, and other recursive elements of a formal sign language

Two lines of evidence point toward the mechanism of dreams that does not depend on the LPFC. First, neuroimaging observations indicate that the LPFC is inactive during the sleep, both slow-wave sleep and REM sleep ^123^. Second, in people whose LPFC gets damaged, dreams do not change at all ^124^. In dreams and hallucinations, images of people and objects may arise spontaneously and even hybridize into novel mental images. A subject can then describe these novel images and nonpresent people verbally, all without ever relying on LPFC-controlled mental synthesis.

We conclude that it is often impossible for an external observer to reliably infer internal imagery of the speaker based on expressive language. The speech may be automatic and independent of internal imagery, triggered spontaneously or driven by sensory input. Even when words reflect internal images, they may not inform of the LPFC control over imagination since even the novel images may have formed spontaneously as in dreaming or hallucination. There is no accurate method for external assessment of mental synthesis with respect to expressive language. While understanding of flexible syntax commonly requires LPFC-controlled mental synthesis, expressive language does not.

Several other bottom-up processes are often erroneously assumed to be indicative of the LPFC-controlled mental synthesis:
1. **Object recognition / perceptual closure.** Recognition of pictures with incomplete visual information is often erroneously assumed to be under the control of the LPFC. While there is no doubt that objects encoded in the posterior cortex can be primed by a category encoded in the PFC, there is strong evidence of bottom-up-driven PFC-independent process in perceptual closure tests, such as Moony faces test ^125^. Accordingly, a good performance at these tests has no bearing on the LPFC-controlled mental synthesis abilities.
2. **Drawing interpretation.** The tendency to perceive objects from lines and colors has been used extensively in psychological tests such as blot test. It is often as easy to recognize an object from meaningless colors or scribbles of lines as it is from a cloud in the sky. Furthermore, once we recognize an object in a drawing, we tend to presume knowing the inclinations of the artist's mind. However, the artist may have produced the drawing automatically or just spilled some colors as in blot test pictures.

Even some representational drawings can be fully automatic. For example, instrumental training (a variation of Pavlovian training) has been used to train elephants to produce representational works of art. Patients with visual agnosia can copy a drawing without any awareness of the type of object in the picture. The famous patient known in literature as pilot John faithfully copied a representation of St. Paul Cathedral without knowing the object in the picture ^126^. Scribbling can often produce recognizable forms with no intention on the part of the scribbler. These examples are reminiscent of the problems with interpretation of expressive language. Outside of a complex dialog, neither expressive language, nor object recognition or drawing can inform on the speaker or artist's LPFC-controlled mental synthesis ability. Only a complex syntactic dialog or representational drawing directed by a verbal description of a *novel object* (“a five headed horse”) can be a reliable indicator of the LPFC ability to combine disparate neuronal ensembles.

## Conclusions

It is commonly believed that there is a strong association between early language acquisition and normal cognitive development (critical period hypothesis) ^2–5^; however, there is no consensus on the neurological mechanism of this association. We hypothesized that the use of infinite syntactic language in childhood, but not of a finite homesign communication system, provides essential experience for developing synchronous connections between the LPFC and the posterior cortex and that those connections are crucial for mental synthesis of disparate objects and understanding of syntactic language. Accordingly, we predicted that without childhood exposure to infinite syntactic language children would not acquire mental synthesis. We tested this hypothesis by analyzing all linguistic isolates reported in scientific literature. Our analysis confirmed that individuals linguistically deprived until puberty performed poorly in all tests of *mental synthesis* despite a focused multi-year rehabilitation efforts. The consistent observation of *mental synthesis disability* in these individuals stands in stark contrast to their performance on memory as well as semantic tests: they could easily remember hundreds of newly learned words and recall previously seen images from memory but had real difficulty in any tasks requiring them to combine these images into novel configurations.

As is the case with ontogenetic development of many other neurological systems from muscle innervation to the development of sensory systems, nature's intent must be complemented by adequate nurture. What is highly unusual about the ontogenetic acquisition of *mental synthesis* is that the necessary experience is provided by the exposure to a purely **cultural phenomenon**: a syntactic language. For the normal development of vision, light reflected from surrounding objects has to reach the retina, but that occurs whenever it is light, independent of cultural exposure; for the normal development of the muscular system, the trophic factors released by muscles have to reach their neurons, but that occurs whenever a child is moving – the stimulation to neurons comes naturally even when a child is growing alone in a forest ^87^. However, this is not the case with mental synthesis. The development of mental synthesis requires a community of humans willing to engage a child with the use of an infinite syntactic language complete with spatial prepositions, verb tenses and other recursive elements; finite homesign systems do not suffice. Furthermore the exposure to an infinite syntactic language has to occur during the period of neural plasticity, which peaks before the age of two ^18^ and expires some time before puberty. These two observations have a profound consequence for education of children, both with or without learning disabilities, but would be especially instrumental for the former, such as those diagnosed with ASD. Children must be encouraged to actively synthesize novel mental images and they have to be provided with tools to facilitate this process: infinite syntactic language, toys to encourage pretend play, drawing, puzzles, and arithmetic exercises. While reading aloud a book of fairy tales stimulates the child's LPFC-controlled mental synthesis (imagining the adventures of cyclops and mermaids), watching TV simply imposes mental images on the child's visual system and therefore is not expected to stimulate LPFC-controlled mental synthesis. With this understanding, future therapeutic modalities targeting mental synthesis may benefit children delayed in development of language ability.

## Acknowledgments

We wish to thank Dr. Petr Ilyinskii for productive discussion and scrupulous editing of this manuscript and Dr. Irene Piryatinsky for help with interpretation of IQ tests.

